# Secreted reporter assay enables quantitative and longitudinal monitoring of neuronal activity

**DOI:** 10.1101/2021.01.22.427811

**Authors:** Ana C. Santos, Sungjin Park

## Abstract

The ability to measure changes in neuronal activity in a quantifiable and precise manner is of fundamental importance to understand neuron development and function. Repeated monitoring of neuronal activity of the same population of neurons over several days is challenging and, typically, low-throughput. Here, we describe a new biochemical reporter assay that allows for repeated measurements of neuronal activity in a cell type-specific manner. We coupled activity-dependent elements from the *Arc/Arg3.1* gene with a secreted reporter, *Gaussia* luciferase, to quantify neuronal activity without sacrificing the neurons. The reporter predominantly senses calcium and NMDA receptor-dependent activity. By repeatedly measuring the accumulation of the reporter in cell media, we can profile the developmental dynamics of neuronal activity in cultured neurons from male and female mice. The assay also allows for longitudinal analysis of pharmacological treatments, thus distinguishing acute from delayed responses. Moreover, conditional expression of the reporter allows for monitoring cell type-specific changes. This simple, quantitative, cost-effective, automatable, and cell type-specific activity reporter is a valuable tool to study the development of neuronal activity in normal and disease-model conditions, and to identify small molecules or protein factors that selectively modulate the activity of a specific population of neurons.

**Significance:** Neurological and neurodevelopmental disorders are prevalent worldwide. Despite significant advances in our understanding of synapse formation and function, developing effective therapeutics remains challenging, in part due to the lack of simple and robust high-throughput screening assays of neuronal activity. Here, we describe a simple biochemical assay that allows for repeated measurements of neuronal activity in a cell type-specific manner. Thus filling the need for assays amenable to longitudinal studies, such as those related to neural development. Other advantages include its simple and quantitative nature, logitudinal profiling, cell type-specificity, and being multiplexed with other invasive techniques.

## Introduction

Neuronal activity plays fundamental roles in the formation and function of neuronal circuits, from synapse development to regulation of synaptic strength to learning and memory (Katz and Shatz, 1996; Flavell and Greenberg, 2008; Sahores and Salinas, 2011; Andreae and Burrone, 2014; Araya et al., 2014; Pan and Monje, 2020). Misregulation of these processes can cause neurodevelopmental, neurological and psychiatric disorders including intellectual disability, epilepsy and schizophrenia (Zoghbi, 2003; van Spronsen and Hoogenraad, 2010; Melom and Littleton, 2011; Zoghbi and Bear, 2012; Lepeta et al., 2016). Despite advances in our knowledge of synapse biology in both normal and disease conditions, developing effective therapeutics remains challenging in part due to the lack of simple and robust high-throughput screening assays of neuronal activity.

In recent years, there has been increased use of new technologies yielding rich and sizeable dataset results, such as RNAseq studies, proteomics and genomics (Geschwind and Konopka, 2009; Wang et al., 2009; Manzoni et al., 2018). Although these studies are typically not designed to identify new drug therapies, they may lead to the unbiased identification of novel drug targets, if coupled with a secondary functional screening platform. In addition to these omics approaches, there has been a concurrent increase in the development of genetically encoded neuronal activity reporters (Tian et al., 2012; Abdelfattah et al., 2019). These tools have been incredibly valuable for the dissection of neural circuits *in vivo*, but are associated with a number of caveats that make them inadequate for the identification of new therapeutic drugs. For example, these are often based on imaging of a fluorescent sensor, which over time can lead to photobleaching, phototoxicity or buffering issues (Bootman et al., 2018; McMahon and Jackson, 2018). This can be problematic because pharmachological and chemico-genetic manipulations can have long-term effects different from their initial response (Ghezzi and Atkinson, 2011; Soumier and Sibille, 2014; Dennys et al., 2015). Thus, monitoring both the acute and long-term effects of pharmacological manipulations on the same population of neurons is critical to developing therapeutics with the intended long-term effects while also minimizing undesired effects, such as drug tolerance.

Here, we developed a novel non-invasive live-cell assay that enables the monitoring of neuronal activity over prolonged periods of time, ranging from minutes to days. Activity can be assayed multiple times because we combined an activity-dependent driver, based on *Arc/Arg3.1* regulatory elements, with a secreted reporter, *Gaussia* luciferase. We show that longitudinal monitoring of the accumulated reporter in culture media reveals the developmental dynamics of neuronal activity in different conditions. Further, direct comparison of changes in neuronal activity within the same population of neurons upon pharmacological manipulation can easily account for the natural variability that exists from culture to culture. Because the reporter is amenable to repeated measurements, temporal analyses can be performed, which allow us to distinguish acute and delayed neuronal responses to different types of perturbations. Furthermore, expression of the reporter in a Cre-dependent manner allows for selective monitoring of neuronal activity in a subpopulation of neurons within heterogeneous cultures. This simple, quantitative, and selective activity reporter is a useful tool to study the development of neuronal activity in normal and disease conditions and to identify factors that selectively modulate neuronal activity.

## Materials and Methods

### Animals

All animal care and experiments were conducted in accordance with NIH guidelines and the University of Utah IACUC committee (protocol no. 18-02004). C57Bl6/J mouse lines were maintained under normal housing conditions with food and water available *ad libitum* and 12h light/dark cycle in a dedicated facility at the University of Utah. All experiments were performed using wild-type mice of either sex.

### Experimental Design and Statistical Analysis

All pair-wise comparisons were analyzed by two-tailed Student’s t-test. We used a one-way ANOVA followed by Tukey multiple comparisons with multiplicity adjusted p-values wherever more than two conditions were tested (Graphpad Prism). Nested statistics were performed whenever possible. For uniformity, all data are plotted as the mean of the total n with standard error of the mean (SEM) error bars. For all experiments, the n numbers shown refer to the number of samples per condition, while N numbers refer to the number of experiments. No statistical methods were used to pre-determine sample sizes, but our sample sizes were similar to those generally employed in the field. Occasionally, control samples are presented in multiple figures, and this is noted in the figure legends. A p value < 0.05 was considered significant, for all tests * p<.05, **p<.01, ***p<.001, ****p<.0001.

### Cell Culture

*Neuronal cultures* were prepared from newborn (P0) wild-type mice using a commonly used protocol. Briefly, hippocampi or forebrain were dissected in HEPES-buffered saline solution (HBSS), enzymatically (using 165 units of papain, Worthington LS003126) and mechanically digested until a single cell suspension is obtained. Cells were then plated in poly-L-lysine (Sigma P2636, 0.2 mg/mL in 0.1M Tris) coated 12-well plates or coverslips. To generate neuron-enriched cultures, cells were treated with AraC (2.5μM, Sigma C6645) after 1 day *in vitro* (DIV) to prevent the proliferation of mitotic cells. Otherwise, AraC treatment was performed on DIV4. Approximately half of the media was replaced with fresh neuronal media every 3 days to prevent evaporation and accumulation of metabolic byproducts. Neuronal media consists of Neurobasal A (Gibco 10888) containing 1% Hyclone horse serum (Fisher Scientific SH30074.03), 1% Glutamax (Gibco 35050), 2% B-27 (Gibco 17504), and 1% Penicillin/Streptomycin (Gibco 15140). In experiments where we test the effect of astrocyte-conditioned media (ACM), neuronal media without serum was used for ½ media change at DIV4 and ACM treatment started on DIV7. ACM was prepared by incubating neuronal media without serum in freshly prepared confluent astrocyte cultures for 24hrs. Neuronal cultures were collected at DIV14-16 for immunostaining and all pharmacology experiments performed at DIV13-15.

*Astrocyte cultures* were prepared from wild-type mice aged P2 using the traditional method. Briefly, the forebrain was dissected, enzymatically and mechanically digested, until a single cell suspension is obtained, as performed for neuron cultures. Cells were plated in poly-D-lysine (50 μg/ml, Millipore A-003-E) coated flasks and grown in astrocyte growth media containing DMEM (Gibco 11960044), 10% fetal bovine serum (FBS, Gibco 16140071), 1% Penicillin/Streptomycin (Gibco 10378016), 1% GlutaMAX (Gibco 35050061), 1% Sodium Pyruvate (Gibco 11360070), 0.5 μg/ml Insulin (Sigma I6634), 5 μg/ml NAC (Sigma A8199), and 10 μM Hydrocortisone (Sigma H0888). The next day a full media change is performed after vigorous shaking and every 2-3 days thereafter, as needed for neuron cultures. If other cell types persisted, these were removed by shaking.

*HEK293T cells* were cultured in DMEM containing 10% FBS, 1% sodium pyruvate, and 1% Penicillin/Streptomycin. Cells were plated at a density of 0.15 million cells per well onto a PEI (25 μg/μl) coated 12-well plate and transfected the next day using the Fugene 6 transfection reagent and according to the manufacturer’s directions (Promega E2691). The same amount of plasmid was transfected per condition. After 24 hrs, media was changed to serum-free DMEM and approximately 24 hrs later both media and lysates were harvested. Media samples were stored short-term at 4C until used for luciferase assays, typically up to one week. HEK293T and Lenti-X 293T lines were not authenticated. All cells were maintained in a humidified incubator at 37°C and 5% CO2.

### Lentivirus Packaging

Lenti-X 293T cells were cultured in DMEM containing 10% FBS, 1% sodium pyruvate and 1% Penicillin/Streptomycin, as described for HEK293T cells. Cells were plated at a density of 2.5-3 million cells per 10 cm dish and transfected the next day using Fugene 6. Packaging plasmids pMD2.G and psPAX2 were obtained from Addgene (plasmids #12259 and 12260, respectively) and used at the ratio 10μg:6μg:10μg (transfer:pMD2.G:psPAX2). 24hours after transfection media was completely replaced and plates returned to the incubator for an additional 48 hrs. Media was collected and filetered through a 0.45 μm PES filter. Lentiviral supernatant was then centrifuged using a benchtop Beckman Optima XP ultracentrifuge at 120,000g for 2 hrs at 4C. The lentiviral pellet was then resuspended in PBS and aliquots stored at −80C until needed. Lentivirus was also commercially packaged (Vigene Biosciences, Rockeville, MD). Transduction was performed by mixing an adequate amount of lentivirus (variable) into neuronal media and incubating this with neurons for 3 hrs, after which a full media change was performed. Cultures were centrifuged at 1000g for 30 minutes to increase transduction efficiency.

### Immunocytochemistry and Imaging

Cells were rinsed with PBS and immediately fixed using 4% paraformaldehyde with 4% sucrose in PBS for 15 minutes. After rinsing with PBS, a 15-minute permeabilization using 0.2% Triton-X100 in PBS, and a 1-hour blocking solution (4% BSA and 4% normal goat serum in PBS), cells were incubated overnight at 4°C in primary antibody diluted in blocking solution (mouse anti-CamKII from Millipore 05-532, rabbit anti-Gluc from Nanolight 401P, mouse anti-GAD67 from Millipore MAB5406, mouse anti-GFAP from Cell Signaling Tech. 3670, chicken anti-MAP2 from Abcam ab5392). After washing with PBS, the cells were incubated with a fluorescent secondary antibody and Hoeschst (Hoechest 33342 from Invitrogen, donkey anti-rabbit Alexa488 from Invitrogen A21206, all others from Jackson ImmunoResearch). Coverslips were mounted onto glass slides using Prolong gold mounting solution (Invitrogen P36930). Imaging was performed using either a Nikon E800 epi-fluorescence microscope or Nikon A1 for confocal imaging.

### Plasmid Cloning

The SNAR construct was synthesized by first introducing a core SARE (cSARE) sequence upstream of the Arc minimal promoter using a long primer. The cSARE sequence is the following: CCTGCGTGGGGAAGCTCCTTGCTGCGT CATGGCTCAGCTATTCTCAGCCTCTCTCCTTTTATGGTGCCGGAAGCAGGCAGGCTGC (85bp). The forward and reverse primers used were: 5’-TACCAAGCTTGGATCCCCTGCACCTGCGTGGGGAAGCTCC TTGCTGCGTCATGGCTCAGCTATTCTCAGCCTCTCTCCTTTTATGGTGCCGGAAGCAGGCAGGCTGC AGATCTCGCGCAGCAGAGCAC-3’ and 5’-CATGGTGGCTGGATCCCTGGTCGTCGGTGCTGCGGCT-3’. The cSARE sequence was derived from the mouse sequence and it is largely conserved in the macaque and human (Kawashima et al., 2009). This PCR product was then introduced into an AAV vector backbone with the humanized *Gaussia* luciferase coding sequence (Tannous et al., 2005) using inFusion cloning. Each cSARE sequence was then synthesized by PCR using this initial construct as a template (the forward and reverse primers used were: 5’-AGCCCCGGGACGCGTAGCCTGCCTGCGTGGGGAAG -3’ and 5’-CAGACTGCAGCCTGCCTGCTT-3’). Three additional cSARE sequences were then added, one at a time preceding the first cSARE. The entire 4x cSARE-ArcMin-Gluc was then cloned as a single insert into an FCK vector (addgene #51694) using restriction enzymes *PacI* and *EcoRI* for use as a transfer plasmid and packaging into lentivirus particles. Secreted Nanoluciferase (sNluc) was obtained by adding the Ig-kappa signal peptide to its N-terminus using a long primer. The primers used were: 5’-AGCTCG CCATGGGCCACCATGGAGACAGACACACTCCTGCTATGGGTACTGCTGCTCTGGGTTCCAGGTTCCACTG GTGAC ACTAGT TATCCATATGATGTTCCAGATTATGCT GGTGGATCA GTCTTCACACTCGAAGATTTCG-3’ and 5’-CGAGCTGAGCTCTTACGCCAGAATGCGTTCGCA -3’, forward and reverse, respectively. sNluc (from addgene plasmid #66579) was then inserted into an FCK vector with the hPGK promoter. The hPGK promoter had been cloned into FCK vector from another addgene plasmid (addgene #74444). Each construct was then cloned into a double-floxed inverted open reading frame (DIO) vector backbone (addgene #87168) to obtain a Cre-dependent expression construct. Floxed constructs were then inserted back into the same FCK vector for lentivirus packaging. The constructs were verified by sanger sequencing.

### Pharmacology

Neurons were allowed to develop for 13 days before any pharmacological agents were applied to the cultures. Tetrodotoxin (TTX), APV, and CNQX were purchased from abcam (cat. no. ab120055, ab120055, ab120044, respectively), Nifedipine from Tocris (cat no. 1075), BAPTA-AM from Invitrogen (cat no. B1207), U0126 from Cell Signaling (cat no 9903), and BDNF from PeproTech (cat. no. 450-02). All others were from Sigma-Aldrich. For all experiments the intial time point coincides with the start of treatment. Typically, SNAR accumulation was measured by calculating the difference from the 16 to 40hr time-points for a 24 hr interval, and normalizing to the initial time-point (0hr). All pharmacology was performed in this manner, except for inhibitor washout experiments. Only experiments where a full media change (washout), was performed the SNAR signal was normalized to the sNluc control.

### Luciferase Assays

To determine luciferase activity, we measured luminescence from media samples upon addition of substrate. Samples, typically 10-20 μl of conditioned media, were loaded onto a 96-well opaque white plate (VWR cat. no. 82050-736). Substrates, Coelenterazine native (CTZ, NanoLight technology cat. no. 303) and Furimazine (FMZ, Promega cat. no. N1110) were added using a micro-injector connected to the plate reader (BioTek Synergy HT). To prevent signal overlap or interactions between substrates, we run CTZ and FMZ reactions separately. CTZ was dissolved in acidic ethanol (0.06N HCl) to a stock concentration of 23.6mM and diluted in 0.1M Tris-HCl pH7.5 before use to a working concentration of 60 μM. FMZ and NanoGlo buffer were purchased from Promega and diluted 4:1 in PBS just before use. 100 μl of substrate were injected per reaction and luminescence signal recorded. Luminescence in CTZ was calculated using the sum of the first 10 seconds of luminescence immediately after injection of substrate, and luminescence in FMZ using an average of 10 readings after a 3-5 min incubation with substrate. Since the luminescence of sNluc in CTZ is linearly proportional to the amount of sNluc in the sample, we can calculate the contribution of sNluc to the CTZ signal by multiplying the FMZ signal of the sample by the constant ratio, *c*, where *c* is the ratio between the luminescence of a sNluc only sample in CTZ and FMZ reactions (*c* = CTZ_sNluc_ /FMZ_sNluc_). By simply subtracting the contribution of sNluc from the total CTZ signal, we can calculate the Gluc signal in the CTZ reaction (Gluc_Sample_ = CTZ_Sample_ – *c* x FMZ_Sample_). Luciferase readings take approximately one hour per 96-well plate and calculations on exported data are simple, as outlined above. Overall, once samples are collected running and analyzing the data of a luciferase assay is relatively fast and results can be obtained within hours.

## Results

### Secreted Neuronal Activity Reporter (SNAR) is a dual luciferase live-cell assay

Expression of the immediate early gene *Arc/Arg3.1* is rapidly induced in response to various stimulation paradigms both *in vitro* and *in vivo* (Lyford et al., 1995; Gouty-Colomer et al., 2016). Enhancer elements of the immediate early gene *Arc/Arg3.1*, namely the synaptic activity responsive element (SARE), have recently been exploited as an activity-dependent driver (Kawashima et al., 2013; Das et al., 2018; Wu et al., 2018). To decrease the size of the reporter we used the conserved core domain of SARE (cSARE, see Materials and Methods for sequence) and generated an activity-dependent driver, which consists of four tandem repeats of cSARE followed by the Arc minimal promoter (Kawashima et al., 2009). We then combined this activity-dependent driver with *Gaussia* luciferase (Fig. 1A). We named this construct Secreted Neuronal Activity Reporter (SNAR). The term neuronal activity is thus used broadly here, since induction of *Arc/Arg3.1* remains topic of research and is not only associated with specific patterns of neuronal firing but also with synaptic plasticity (Korb and Finkbeiner, 2011). To generate a control reporter that could be used simultaneously, Nanoluciferase was coupled with the human PGK promoter to make a constitutively secreted control. Both SNAR and control constructs were designed to be compact (1.3kbp and 1.2kbp, respectively) so that they can be efficiently delivered into neurons using a variety of approaches, including AAV and lentivirus (lenti).

**Figure 1.**
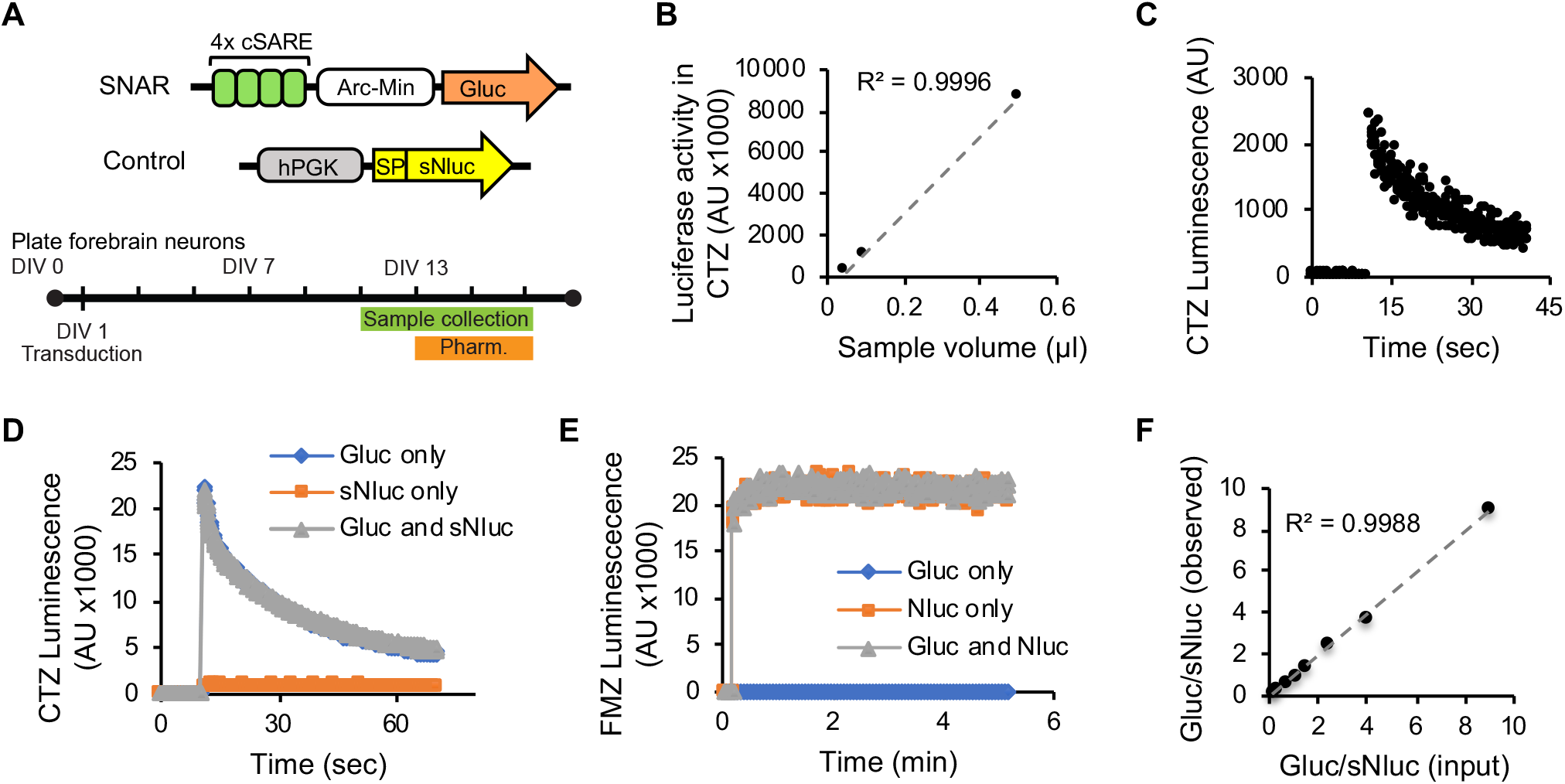
Dual Secreted Luciferase Assay. Gluc and sNluc are sensitive reporters and can be independently measured from mixed samples. **A.** Diagram of activity-dependent (SNAR), and control (hPGK) constructs, and a typical experimental paradigm. For most pharmacology experiments drugs were applied on DIV13 and the accumulation of reporter in the media was monitored over the following 48hrs. **B-C.** SNAR was transduced into primary neurons and allowed to accumulate in culture media for 10 days. **B.** SNAR activity from several dilutions of a medium sample. **C.** An example trace of Gluc kinetics of 0.05μl of the medium sample used in *B.* **D-F.** HEK293T cells were transfected with either a CAG_Gluc or CAG_sNluc plasmid and conditioned media collected. **D-E.** Kinetics of Gluc (D) and sNluc (E) are not affected in mixed samples from HEK293T conditioned media. Representative kinetic plots are shown for luciferase reactions in CTZ (D) and FMZ (E). **F.** Mixtures of both luciferases were prepared in various ratios (input ratio) and the activity of each luciferase was measured. These results were then used to calculate the Gluc/sNluc ratio (observed ratio). The observed ratio accurately reflects the input ratio (slope = 0.9944, *R²* = 0.9988).

We chose to use *Gaussia* Luciferase (Gluc) and Nanoluciferase (Nluc) in the SNAR assay for their small size, and superior brightness (Shao and Bock, 2008), as these characteristics would impart increased sensitivity to the assay. Indeed we found that Gluc can be detected in a sub-microliter volume of medium (Fig.1B, C) Importantly, Gluc is endogenously secreted affording us the ability to perform longitudinal studies since cell lysis is not required (Suzuki et al., 2007). Similarly, Nluc can easily be engineered to become secreted using a signal peptide (Hall et al., 2012; England et al., 2016). Previous studies reported that Gluc and Nluc have distinct kinetics and substrate specificity and can be combined as a dual luciferase system (Heise et al., 2013; Wires et al., 2017).

To determine whether we could apply this system to the SNAR assay, we started by validating that Gluc and secreted Nluc (sNluc) could be independently measured from culture media. We transfected HEK293T cells with Gluc or sNluc under a constitutive promoter (CAG) and collected media samples over time (up to 6 hours). We found that both Gluc and sNluc were secreted and linearly accumulated in the culture medium over time (not shown). The kinetic properties of each luciferase were not affected by the other in a mixed sample (Fig. 1D, E). Furimazine (FMZ), is a specific substrate of sNluc and does not react with Gluc. Coelenterazine (CTZ), a robust substrate for Gluc, also reacts with sNluc, albeit at very low level (Fig. 1D). Since the contribution of sNluc in CTZ signal is linearly propotional to the FMZ signal in a mixed smaple, we were able to reliably calculate the Gluc/Nluc ratio from mixed samples and found it to match the input Gluc/Nluc ratio (see Materials and Methods for formula. Fig. 1F, slope = 0.9944, *R²* = 0.9988). These results show that Gluc and sNluc can be used reliably in a secreted dual luciferase assay, making them ideal for use in the SNAR reporter.

### SNAR reflects neuronal activity

To determine whether SNAR could reliably measure neuronal activity, we tested if manipulating neuronal activity would lead to changes in reporter accumulation in the culture medium. Mouse primary cortical neurons were infected with lenti:SNAR and lenti:Control on DIV1, allowed to mature, and treated with pharmacological reagents on DIV13. SNAR activity was then monitored over the following 2 days from DIV13 to DIV15 (Fig. 1A). To inhibit neuronal activity, we treated neurons with a cocktail of inhibitors (TAC: TTX, APV, and CNQX, inhibitors of voltage-gated sodium channels, NMDA receptors, and AMPA receptors, respectively). We started to detect a reduction in reporter accumulation approximately 16 hours after the inhibitors were added and the rate of reporter accumulation in the medium dramatically declined during the following 24 hours (Fig. 2A). In control conditions, however, basal reporter accumulation continues to linearly increase in the medium. This delayed response is likely due to the ongoing release of pre-synthesized protein in the secretory pathway and continued protein synthesis from pre-existing transcripts. Conversely, stimulation of network activity by picrotoxin (PTX, a GABA_A_ receptor inhibitor), robustly enhanced SNAR activity (Fig. 2A, B). In addition, acute stimulation of neurons by washout of inhibitors rapidly induced SNAR activity within 30 minutes (Fig. 2C, D). Notably, temporal analyses also show the rate of SNAR accumulation returns to its basal rate after 2 hours, consistent with the previously reported transient dynamics of *Arc/Arg3.1* expression (Cole et al., 1989; Bramham et al., 2008; Steward et al., 2017; Das et al., 2018). Further, the magnitude of SNAR increase after 30 minutes of inhibitor washout (Fig. 2D, ~4 fold) is similar to that of Arc mRNA as observed in similar experiments performed by other groups (Das et al., 2018). Overall, these results suggest that SNAR can be used to monitor changes in neuronal activity in live neurons and reveal the temporal dynamics of neuronal responses within the same neuronal population.

**Figure 2.**
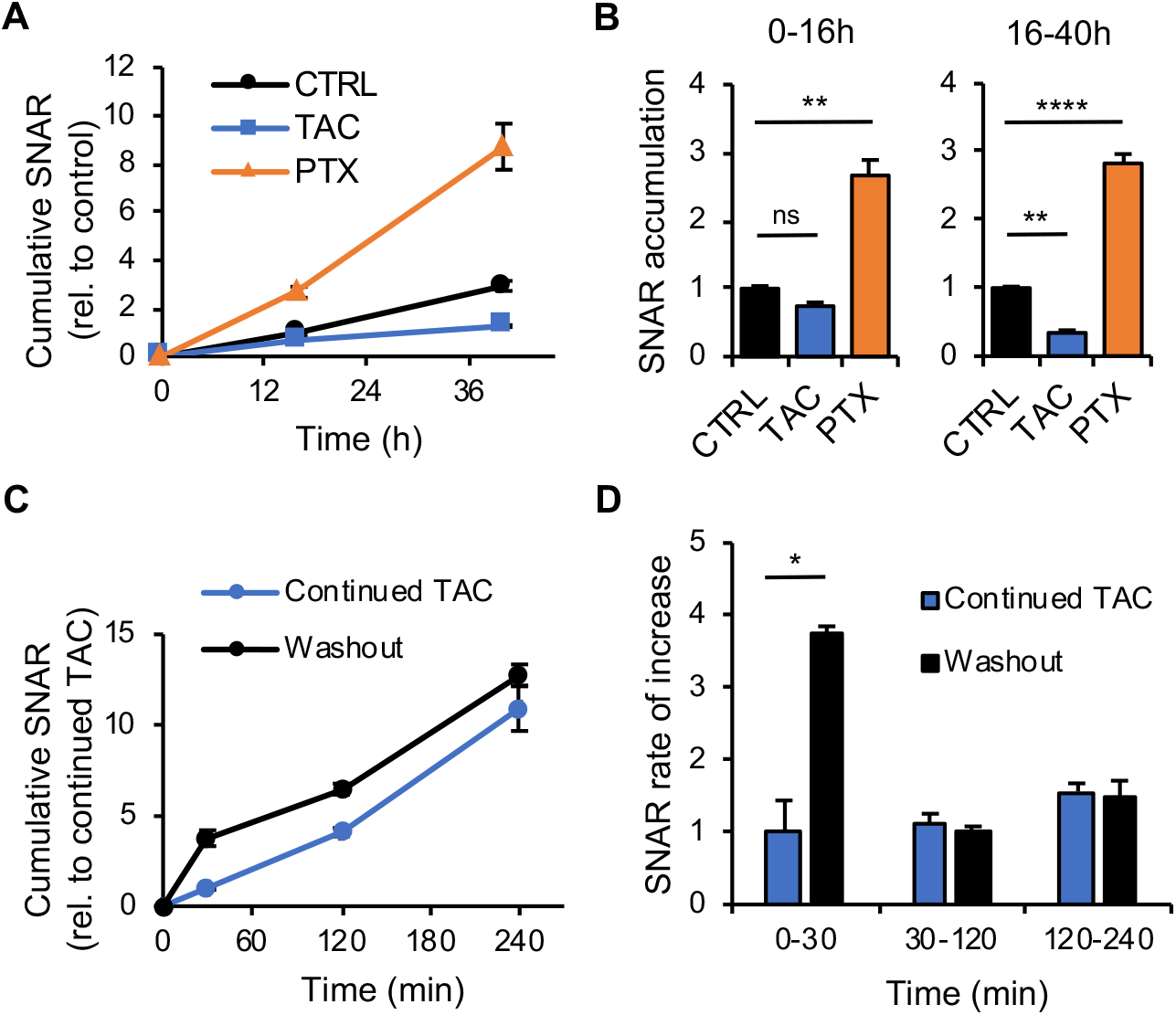
SNAR is an activity-dependent assay. **A.** Primary mouse neuron cultures were treated with an inhibitor cocktail (TAC: 2μM TTX, 50μM APV, and 10μM CNQX), picrotoxin (PTX, 50μM) or a vehicle control for 40hrs total. No media change was performed. Media samples were collected at different timepoints starting at the time of treatment. Cumulative SNAR is normalized to the initial time point (0hr) and relative to 16h control. **B.** Quantification from *A*. SNAR accumulation was measured by calculating the difference from 0 to 16hr and 16 to 40hr time-points, and normalized to the initial time-point (0hr) relative to control. (Nested one-way ANOVA, N=3, n=23 Ctrl, n=18 TAC, and n=11 PTX, ** p=0.001 TAC 0-16h and p=0.084 at 16-40h, **** p<0.0001). **C**. Washout of inhibitors (TAC) induces a rapid increase in SNAR. Neurons were pre-treated with TAC for 48hrs, followed by a full media change to remove accumulated SNAR and replaced with fresh media containing inhibitors (Continued TAC) or no inhibitors (Washout). Cumulative SNAR is normalized to sNluc control and relative to 30 min continued TAC. **D.** Quantification from *C*. Rate of increase is calculated as the change in SNAR in each time interval per unit of time normalized to sNluc control relative to continued TAC (Nested t-test p=.0485, N=3, n=12). *Gaussia* Luciferase (Gluc), secreted Nanoluciferase (sNluc).

### Pharmacological and pathway analysis of SNAR

Inhibition of network activity by an inhibitor cocktail suppressed SNAR activity (Fig. 2A, B). To investigate the specific signaling pathways regulating SNAR, we treated neurons with individual inhibitors. Consistent with previous studies on the endogenous *Arc/Arg3.1* gene and SARE reporter (Rao et al., 2006; Kawashima et al., 2009; Kawashima et al., 2013), inhibition of NMDAR-mediated transmission by APV or blocking neuronal firing by TTX dramatically suppressed SNAR activity (Fig. 3A). Treatment with both TTX and APV showed similar inhibition of SNAR activity as APV alone. Interestingly, blocking AMPAR-mediated transmission with CNQX induced a robust increase in SNAR accumulation between 16-40hours after treatment (Fig. 3A). Although a previous study reported that CNQX paradoxically increases the expression of endogenous *Arc/Arg3.1* mRNA (Rao et al., 2006), the underlying signaling mechanism remains unclear. Long-term inhibition of synaptic activity can elicit homeostatic responses that enhance the efficacy of synaptic transmission and/or neuronal excitability (Turrigiano et al., 1998; Watt et al., 2000). Indeed, the SNAR increase triggered by prolonged CNQX treatment is completely blocked by APV or TTX co-treatment (Fig. 3A). Overall, this assay is useful not only to detect the rapid effects of acute neuronal stimulation (Fig. 2C, D) but also to reveal delayed responses such as due to chronic inactivity (Fig. 3A).

**Figure 3.**
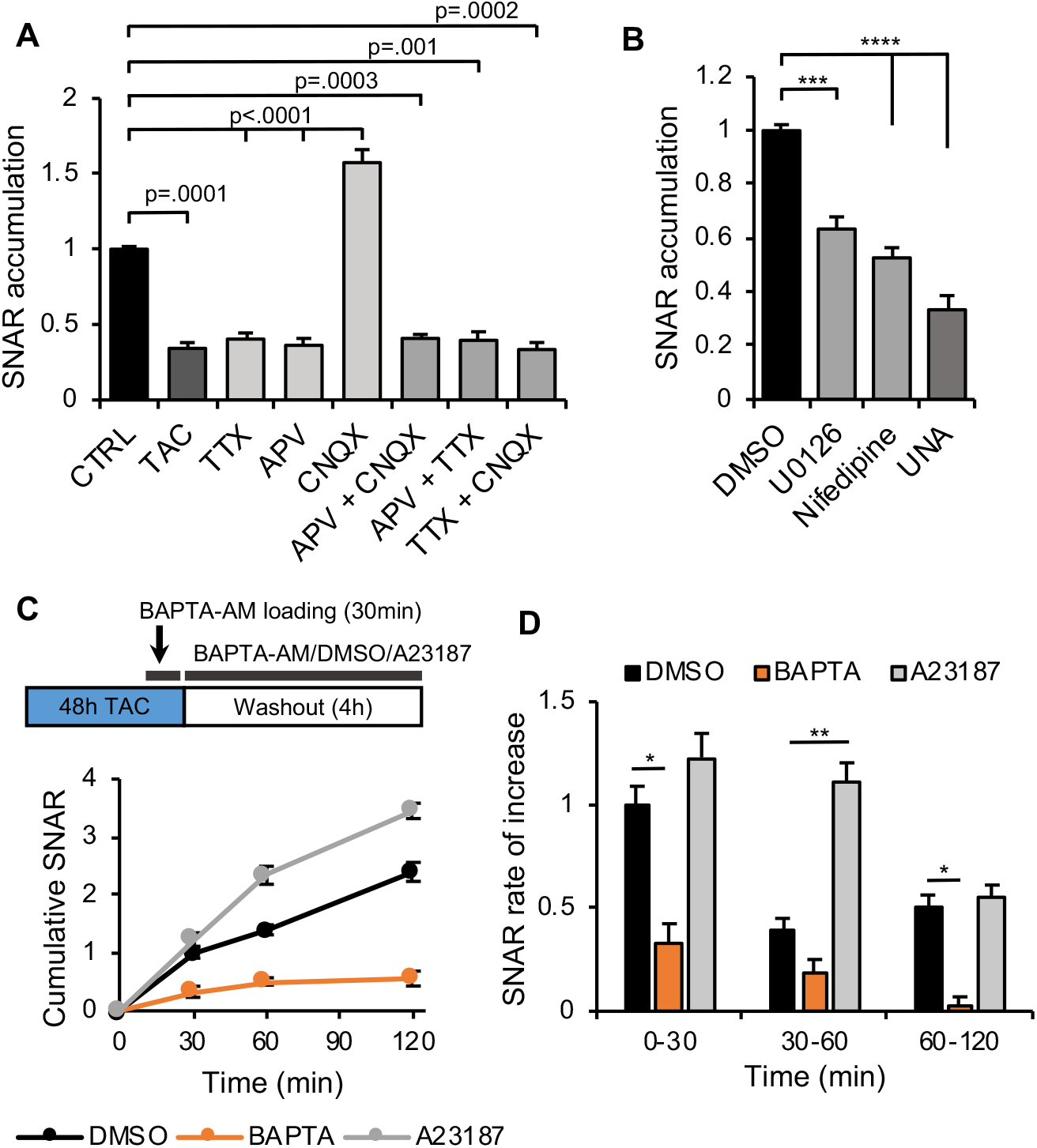
Pharmacological characterization of SNAR induction. **A.** Treatment of WT neurons with various inhibitors shows that SNAR largely reflects NMDAR activity. SNAR accumulation is from a 24hr interval, normalized to the initial time-point (0hr), and relative to control. (N, n): (10, 38) CTRL, (3, 18) TAC, (5, 17) 1μM TTX, (4, 14) 50μM APV, (6, 21) 10μM CNQX, (4, 15) APV and CNQX, Nested ANOVA. Control and TAC conditions from Fig. 2 are replotted here. **B.** ERK signaling and L-type Ca2+ channel activation contribute to SNAR (UNA: 10μM U0126, 10μM Nifedipine and 50μM APV). Same experimental timepoints as in *A.* (N, n): (9, 44) DMSO, (4, 16) U0126, (4, 15) Nifedipine, (3, 11) UNA, Nested ANOVA, *** Tukey p=.0003, **** Tukey p<.0001. **C-D.** BAPTA-AM (25μM) robustly blocks SNAR induction after inhibitor washout, while A23187 (1μM) further increases it. Neurons were treated with TAC inhibitors for 30h preceding washout to allow for detection of bidirectional effects. **C.** Experimental diagram (top). BAPTA-AM was loaded for 20-30 minutes immediately before washout. Cumulative SNAR is normalized to the initial time point (0 min) and a previous timepoint before pre-treatment. **D.** Quantification from *C*. Rate of increase is calculated as the change in SNAR per time interval normalized to the initial time point (0h) relative to control (N=3, n=12, Nested ANOVA, 0-30min Tukey p=.028, 30-60min Tukey p=.0053, 60-120min Tukey p=.04).

Previous studies show that inhibition of ERK1/2 and L-type calcium channels reduces the expression of *Arc/Arg3.1* (Murphy et al., 1991; Waltereit et al., 2001; Kawashima et al., 2009). Consistent with previous observations, U0126 (ERK1/2 inhibitor) and Nifedipine (L-type calcium channel blocker) significantly suppressed SNAR accumulation (Fig. 3B). Combining U0126, Nifedipine, and APV did not further reduce SNAR activity compared with APV alone, indicating that NMDAR-mediated signaling plays a predominant role in SNAR activity (Fig. 3B).

Calcium is a poweful second messenger and many calcium reporters, such as GCaMPs, are widely used to monitor neuronal activity (Tian et al., 2012). Therefore, we next asked whether SNAR was activated in a calcium dependent manner. We inhibited activity for 30h using the inhibitor cocktail TAC and tested whether treatment with the cell permeable calcium chelator BAPTA-AM blocked the induction of SNAR observed by inhibitor washout (Fig. 1D). We observed that BAPTA-AM robustly blocked SNAR expression (Fig. 3C, D). On the other hand, treatment with the calcium ionophore A23187 rapidly induced SNAR expression beyond control levels. Overall, these results indicate that SNAR senses neuronal activity in a calcium dependent maner conducted predominantly by NMDAR-mediated transmission and, to a lesser extent, by voltage-gated calcium channels. It is also sensitive to other signaling cascades such as the MAPK pathway.

### Longitudinal monitoring of neuronal activity

Next, we tested if we could use the reporter to monitor neuronal activity over longer periods of time. This would be a substantial advantage over existing techniques since it would allow us to study the dynamics of biological and pharmacological agents over time. In addition, we would be able to monitor the developmental dynamics of neuronal activity with temporal specificity. Given the sensitivity of *Gaussia* luciferase, we observed that the assay could be performed with a very small volume of media (Fig. 1C). This allows for multiple time points to be collected without significant changes to the culture conditions. For this reason, reporter activity can be normalized to a time just prior to treatment, which allows the detection of changes that may be very small or otherwise difficult to capture.

We first determined the stability of secreted Gluc and sNluc in neuron cultures. We infected primary mouse neuron cultures with lenti:SNAR and lenti:Control and transferred conditioned media from the infected neurons to a non-infected culture on DIV7. Repeated measuring of each reporter in the culture medium of non-infected neurons over several days shows that both Gluc and sNluc are stable over several days in culture medium, indicating that the secreted reporters are not degraded in the medium nor taken up by neurons (Fig. 4A). Thus, the stability of the secreted reporters in the culture medium of live neurons allows for long-term monitoring of the reporters.

**Figure 4.**
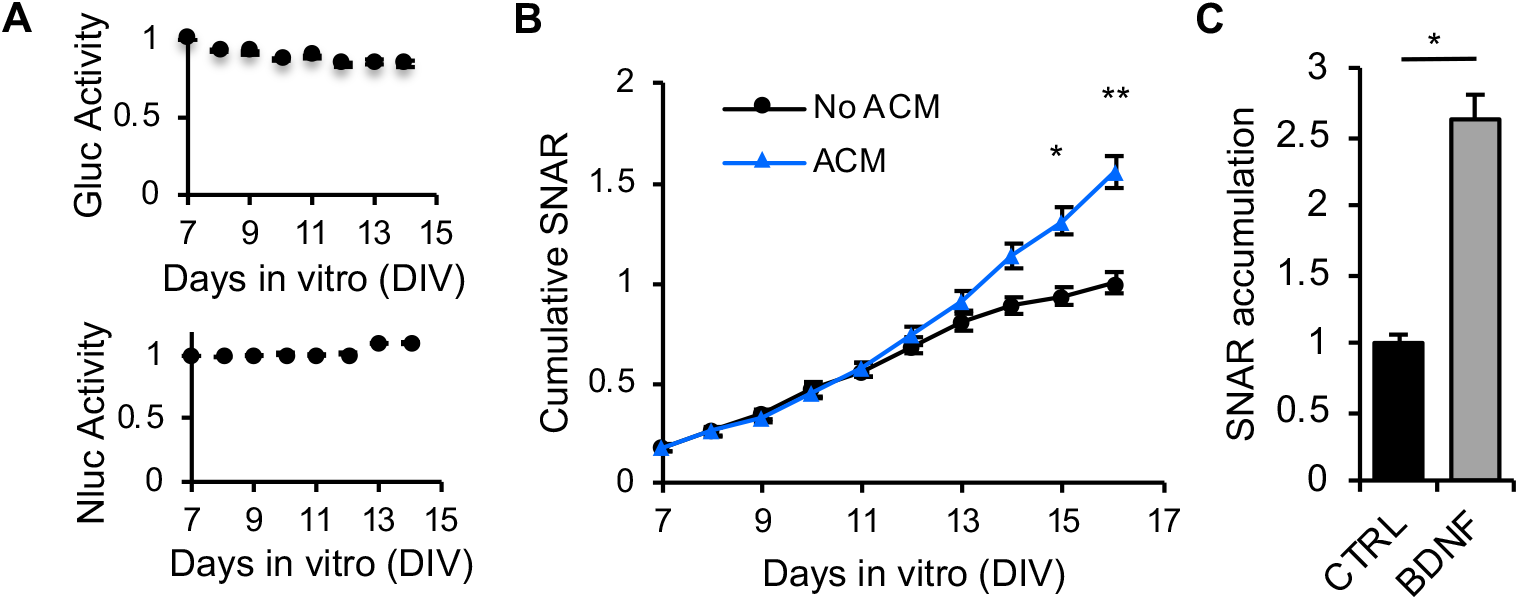
Longitudinal measurement of neuronal activity. **A.** Gluc and sNluc are stable in neuronal media. Neurons were infected at DIV1 and luciferase allowed to accumulate in the media for 7 days, at which point it was transferred to an uninfected neuron culture at the same stage. The activity of each luciferase was monitored for an additional 7 days (N=1, n=5). **B.** ACM induces increased SNAR expression. This increase is most evident after DIV13. WT hippocampal neurons were cultured for a total of 16 days in vitro (DIV). Half the culture medium was replenished with either neuronal media conditioned in astrocytes (ACM) or unconditioned neuronal media (no ACM) on DIV7, DIV10, and DIV13 (N=3, n=12, Nested t-test, *p=.03, **p=.0064). Luciferase accumulation is normalized to end-point value of the control condition. **C.** BDNF treatment (50 ng/ml) at DIV13 robustly increases SNAR expression. SNAR accumulation is from a 24hr interval, normalized to the initial time-point (0hr), and relative to control (N=3, n=25, Nested t-test p=.025).

Astrocytes are important regulators of excitatory synapses and neuronal activity (Chung et al., 2015). It has been demonstrated that astrocytes secrete multiple diffusible factors that promote excitatory synapse formation and function (Araque et al., 2014; Bernardinelli et al., 2014; Allen and Eroglu, 2017; Brancaccio et al., 2017; Blanco-Suarez et al., 2018). Hence, we tested if SNAR could detect the effects of astrocyte-derived factors in synapse development and neuronal activity. We treated neurons with either astrocyte conditioned media (ACM) or unconditioned media (no ACM) and monitored reporter activity daily. In both conditions, we observed a gradual increase in SNAR accumulation, consistent with neuronal maturation and increased synapse number (Chanda et al., 2017). Treatment of neurons with ACM enhanced SNAR activity consistently from DIV12 and the difference increased over the following days (Fig. 4B). These results suggest SNAR can be used to monitor the development profile of neuronal activity *in vitro* and test the activity of non-neuronal factors on synapse development and function.

To determine if SNAR could detect the effects of a single protein factor on neuronal activity, we treated neurons with BDNF, which enhances synapse formation/maturation and transmission via multiple mechanisms (Bamji et al., 2006; Zhou et al., 2006). BDNF treatment at DIV13 significantly and robustly increased reporter activity compared to a vehicle control (Fig. 4C). Overall, the SNAR assay is able to distinguish both acute and delayed effects of pharmacological manipulations and provide mechanistic insights from kinetic analyses.

### Cell type-specific expression

Primary neuron cultures are comprised of heterogeneous neuronal populations. To determine the type of cells expressing SNAR, we performed immunostaining for Gluc together with cell type-specific markers. Consistent with the expression of endogenous *Arc/Arg3.1* (Lyford et al., 1995; Vazdarjanova et al., 2006), the majority (81.33%) of cells expressing SNAR (Gluc-positive) were CamKII-expressing excitatory neurons (Fig. 5A). We also observed that a small population (9.37%) of SNAR-expressing cells were GAD67-positive inhibitory neurons (Fig. 5B). These results demonstrate that SNAR is predominantly expressed in excitatory neurons. Further, to determine whether SNAR expression was specific to neurons and not glia, we prepared a neuron and glia mixed culture. Instead of inhibiting glial proliferation with AraC, we allowed glia to proliferate. As expected, we found that SNAR is not expressed in GFAP-positive astrocytes but it localized to neurons marked by MAP2 (Fig. 5E).

**Figure 5.**
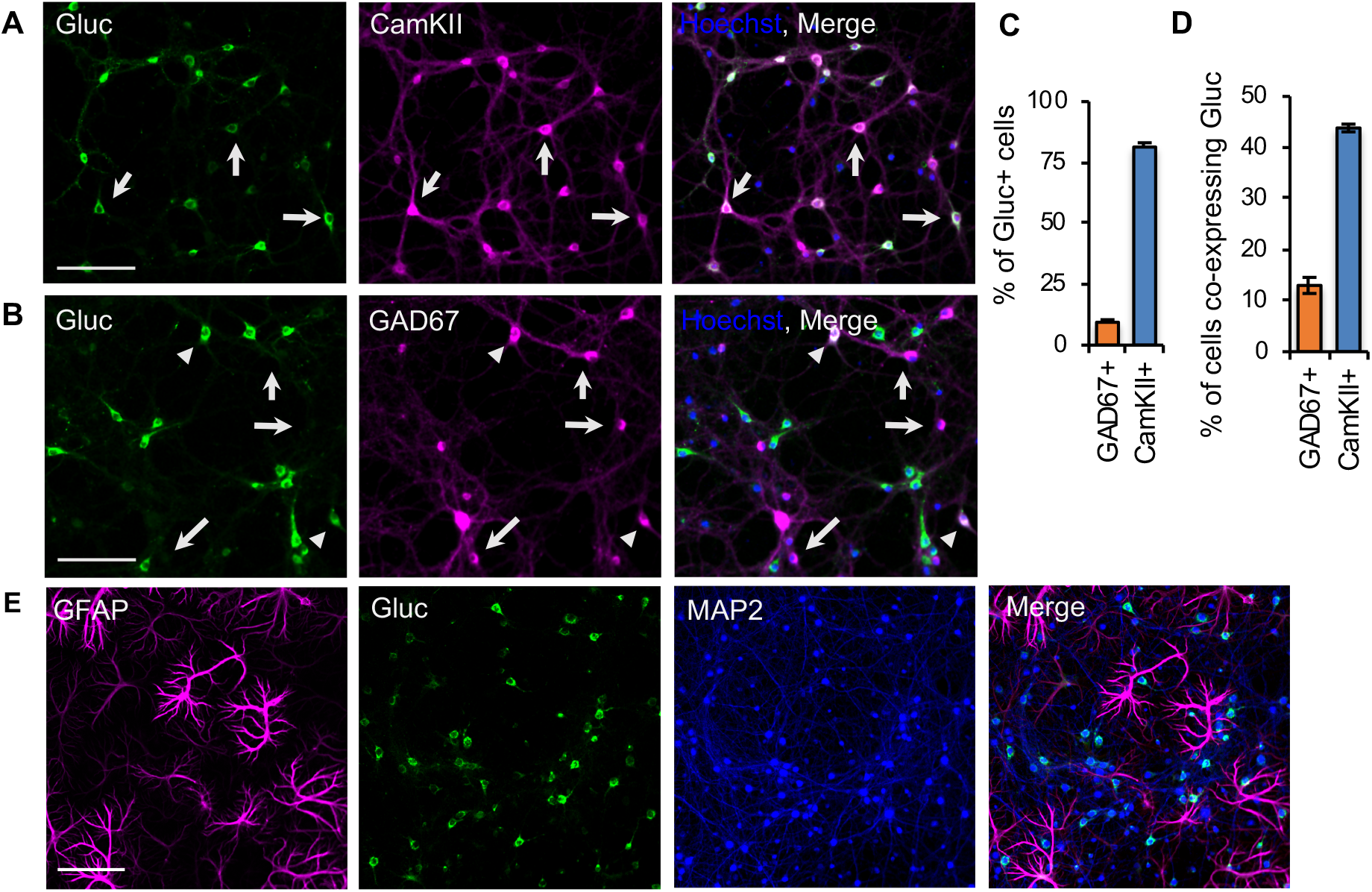
Cell-type specificity of SNAR expression. Immunolabelling of neuron cultures for Gluc-the SNAR reporter, and cell-type specifc markers. Neurons were naïve to any pharmacological treatment until fixation. **A.** Representative images of Gluc and the excitatory neuron marker CaMKII. Arrows indicate Gluc and CaMKII co-expressing cells. **B**. Representative images of Gluc and the inhibitory neuron marker GAD67. The majority of inhibitory neurons do not express SNAR (arrows), however, a small subpopulation does express Gluc (arrowheads). **C.** Characterization of SNAR expressing cells by quantification of Gluc colocalization with CaMKII or GAD67. SNAR is predominantly expressed in CaMKII-positive neurons (81.33%, arrows). Gluc also co-localizes with a small percentage (9.37%) of GAD67+ cells. **D**. Quantification from *A, B*. Most inhibitory neurons do not express SNAR, only 14.5% of GAD67-positve cells co-express Gluc. While 59.8% of CaMKII-positve neurons express clearly detectable levels of Gluc. This could be an underestimate as neurons were not stimulated. **E**. Representative images of Gluc and the astrocyte marker GFAP. SNAR does not co-localize with GFAP, but it does co-localize with MAP2 indicating SNAR expression is specific to neurons and not astrocytes. All scale bars are 100μm.

Having the reporter be expressed in a cell type-specific manner would be particularly advantageous since a number of neurological disorders are caused by cell type-specific defects (Willsey et al., 2013; Zhang and Shen, 2017; Skene et al., 2018). Given the sensitivity of SNAR, which can be detected in a sub-microliter volume of medium (Fig. 1C), we envisioned SNAR would be a useful tool to reveal the subtype-specific response within heterogenous cultures. Therefore, we inserted two lox sites (lox*N* and lox*2272*) flanking the luciferase sequence such that it would only be expressed in the presence of Cre recombinase. To demonstrate that we could combine the Cre system with the SNAR reporter, we transduced neurons with the floxed version of the reporter as well as CamKII:Cre. We found there was very little to no detection of the reporter in the absence of Cre recombinase. However, with Cre expression SNAR was robustly expressed (Fig. 6B). Treatment with the NMDAR blocker, APV, reduced reporter accumulation similar to that of the non-specific reporter (Fig. 6C, 3A). Therefore, by using a cell type-specific promoter to drive expression of Cre, the SNAR reporter can be expressed specifically in a subpopulation of neurons without affecting its activity.

**Figure 6.**
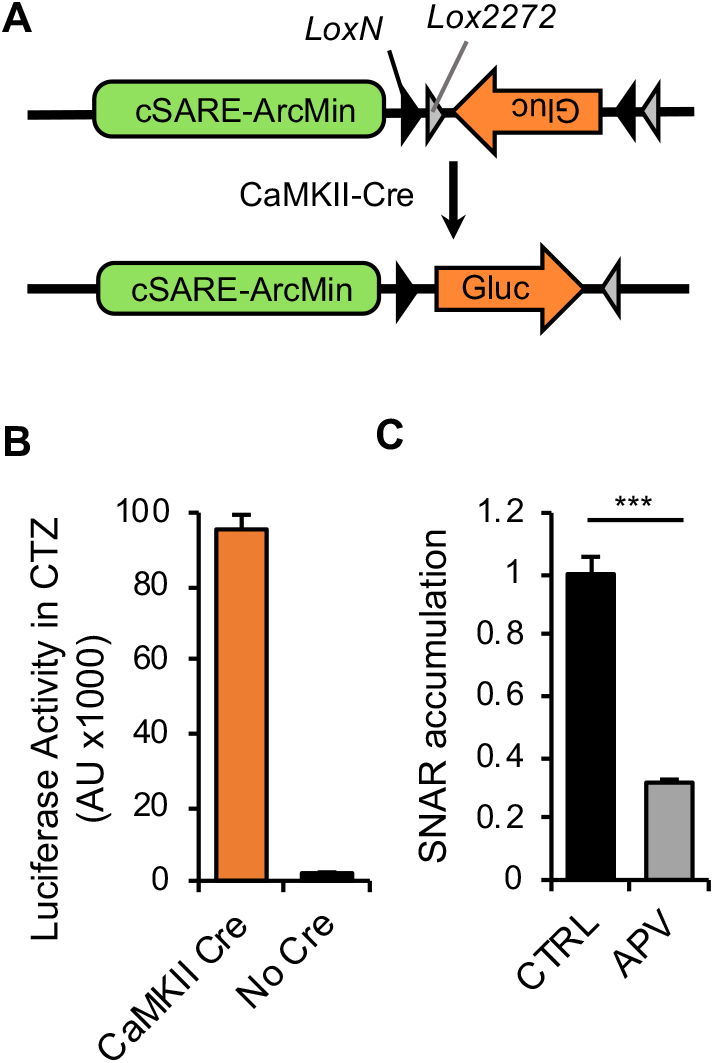
Expression of SNAR in a neuronal subpopulation. **A.** Diagram of the Cre-dependent constructs used. We used a double-floxed inverted open reading frame cassete (DIO). **B.** Neurons transduced with the floxed constructs depicted in *A* express little to no luciferase, while neurons transduced with both CaMKII-Cre and DIO-SNAR show robust Cre recombination and high expression of luciferase. **C**. SNAR expression remains largely mediated by NMDA receptors after Cre recombination. SNAR accumulation from a 24hr interval, normalized to the initial time-point (0hr), and relative to control (N=1, n=4, t-test p=8.33×10^−5^).

## Discussion

Here, we present a novel live-cell assay to quantify changes in neuronal activity over a wide range of time scales. The assay is simple, automatable, and easily performed in a standard molecular biology laboratory. The sensitivity and robustness of SNAR allow for a quantitative assay to be run with a very small volume of culture medium. Therefore, the neuronal activity of the same population of neurons can be measured multiple times and over long periods of time with minimal perturbation to culture conditions. Moreover, neuronal activity itself is susceptible to modulation by other mechanisms such as intrinsic excitability, synaptic plasticity and homeostatic scaling. Thus, monitoring both the acute and long-term effects of pharmacological manipulations on the same population of neurons is critical to developing therapeutics with the intended long-term effects while also minimizing undesired effects.

### Comparison with current methods

Despite the availability of various methods to monitor changes in neuronal activity, the lack of effective therapeutics suggests these are not efficient high-throughput screens. These methods are also inadequate for longitudinal studies, which would be an extremely valuable feature to those studying development an neurodevelopmental disorders and long-term drug effects. For example, current methods to study synapse formation largely rely on immunostaining of fixed neurons and electrophysiological analyses of individual neurons (Basarsky et al., 1994; Christopherson et al., 2005; Ippolito and Eroglu, 2010; Muller et al., 2018; Sudhof, 2018). These methods provide detailed spatial resolution, molecular composition, and mechanistic insights on synaptic transmission of individual neurons. However, they are not ideal to longitudinally monitor activity within a population of neurons because they are endpoint assays. Furthermore, the heterogeneity of cultured neurons creates considerable statistical variability (Wagenaar et al., 2006; Belle et al., 2018). Thus an assay comparing different neuronal populations for different conditions makes it more difficult to identify hits from medium/high-throughput screens.

Alternatively, non-invasive methods have been developed to monitor changes in neuronal activity of the same neuronal population, which include imaging approaches and multielectrode arrays (MEA). Imaging approaches include genetically encoded sensors, such as calcium indicators (Nakai et al., 2001), neurotransmitter sensors (Marvin et al., 2013; Patriarchi et al., 2018), and voltage indicators (Kralj et al., 2011). Although these reporters are ideal for live-cell imaging of network activity (Broussard et al., 2014; Emiliani et al., 2015), fluorescence-based methods are associated with a number of caveats, such as photobleaching, phototoxicity and buffering action (McMahon and Jackson, 2018). MEAs have been used to non-invasively monitor network activity and are useful to identify drugs that affect overall population activity, but cost and non-selectivity limit its application to large-scale screens of neuronal subpopulations (Odawara et al., 2016).

Compared to conventional methods, the SNAR assay provides several advantages for monitoring neuronal activity over time:

#### (1) Kinetics

Because the SNAR assay is designed for multiple time-point measurements, it is suitable to detect changes over a variety of time scales. The SNAR assay can detect changes in activity in as little as 30 minutes of stimulus onset (Fig. 2D) or after multiple days (Fig. 4B). Moreover, kinetic analyses may reveal drug stability, tolerance, or other undesired complications developed over time. This is a major technical advancement since other tools, such as calcium indicators, are not well suited for long-term longitudinal studies.

#### (2) Reduced Variability

By repeatedly monitoring reporter accumulation within the same population of neurons, the effect of drug treatments on reporter activity can be normalized to its accumulation rate prior to the treatment, which significantly lowers variability compared to a conventional assay that compares different neuronal populations for different conditions. This basal accumulation rate also serves as an internal control for variability of culture conditions, including infection rate, neuronal survival, health, and maturation status.

#### (3) Sensitivity and Simplicity

The assay is extremely simple and cost-effective. Luminescence is easily quantified by collecting a small volume of media and mixing it with the respective luciferase substrate, a procedure that can be performed in a conventional laboratory setting and fully automated. Gluc exhibits flash luminescence kinetics upon reacting with a substrate (Fig. 1D), thus a programmable injector is required to consistently measure the initial strong signal. For the plate reader that is not equipped with an injector, a modified Gluc that shows more stable luminescence such as slow-burn Gluc (sbGluc) and superluminescent Gluc (slGluc) would be favored for consistent measurements across multiple samples (Welsh et al., 2009; Gomez-Ramirez et al., 2020).

#### (4) Cell-type Specificity

Unlike MEAs, which are not suitable to cell-type specific analyses, our floxed reporter allows for monitoring of neuronal activity from specific subpopulations of neurons, while maintaining the activity of neighboring and other types of neurons intact. This is, however, contingent on the expression of *Arc/Arg3.1*. For example, SNAR is not expressed in the majority of inhibitory neurons (Fig. 5B).

#### (5) Non-invasive

The use of SNAR is not exclusive. Because we can assay SNAR activity from live-neurons and therefore don’t need to sacrifice them, the reporter can be combined with most other experimental techniques, including immunostaining or electrophysiology.

### Considerations or limitations

#### (1) SNAR activation

Although the SNAR assay provides good temporal resolution, by which stage-specific effects of genetic or pharmacological manipulations can be revealed, it provides limited mechanistic insight. To distinguish whether a specific manipulation alters an early developmental process or directly modulates synaptic transmission *per se*, additional experiments should be performed. In addition, synapse development and function can be affected by non-synaptic effects, such as impaired metabolism and viability of neighboring neurons, therefore, the effects of a hit on the control reporter and cell viability need to be validated. Our results suggest SNAR reflects predominantly NMDAR activity (Fig. 3A). However, to a smaller extent it can also detect the effects of other signaling pathways including the ERK/MAPK pathway. This is not surprising, given *Arc/Arg3.1* plays important roles in multiple pathways, including plasticity mechanisms (Korb and Finkbeiner, 2011). Importantly, we found SNAR signal is highly dependent on cytosolic calcium levels. This is consistent with previous studies on *Arc/Arg3.1* and SARE (Kawashima et al., 2009; Zheng et al., 2009).

#### (2) Temporal dynamics

Due to the on-going release of pre-synthesized protein and newly synthesized protein from pre-transcribed mRNA, there is a lag-time before the reduction in SNAR activity can be detected. We typically measure the changes in SNAR activity after the lag-time (from 16 to 40 hours after stimulus onset). However, acute inhibitory effects can be revealed by quantifying the SNAR accumulation rate after neuronal stimulation. For example, BAPTA acutely blocks the SNAR accumulation induced by inhibitor washout (Fig. 3D). Activation kinetics of SNAR should also be considered when experiments aim at the identification of neuronal activators. Depending on the stimulus, increased neuronal activity can be induced within minutes (Fig. 2D) or gradually over days (Fig. 4B). Thus, both short-term (< 1 hour) and long-term (days) measurements of SNAR activity are recommended to identify spike-like and chronic neuronal activators, respectively. Changes in activity paterns in the order of seconds/minutes might not be detected by SNAR. For example, limited bursts of activity and low tonic activity might not be temporally resolved by SNAR. It should also be noted that SNAR does not have the capability to detect activity in the order of single action potentials, in part due to the distinct time scales of these events.

#### (3) Excitation and inhibition (E/I) balance

E/I balance is tightly controlled and often impaired in disease (Yizhar et al., 2011; Nelson and Valakh, 2015; Selten et al., 2018). However, SNAR does not distinguish whether overall changes in network activity are caused by a direct effect on excitatory neurons or the opposite effect on inhibitory neurons. Moreover, if a drug inhibits both excitatory and inhibitory inputs to the same degree, E/I balance will be maintained and thus may not be detected by the assay and cause false-negative results. These issues can be addressed using the conditional expression of SNAR and/or pharmacology.

#### (4) Spatial Resolution

Although SNAR does not provide specific information pertaining to where the reporter is being expressed because the reporter, *Gaussia* Luciferase, is secreted, the assay can easily be combined with other methods that do provide good spatial resolution, such as immunostaining.

#### (5) Normalization

Since neuronal cultures are typically heterogeneous and sensitive to culture conditions, normalization of the SNAR signal is critical to compare different samples. We observed that normalizing SNAR changes to the pretreatment level of the same neurons produces the most consistent results. However, the control reporter needs to be used when pretreatment levels cannot be measured or neurons are cultured under different conditions from the beginning (as when comparing different genotypes).

### Disease models and other applications

In recent years, human-iPSC derived neurons and brain organoids have become a valuable tool to study development, disease, and human-specific processes as well as to test novel therapeutics (Watanabe et al., 2017; Khan et al., 2020). Although an *in vitro* model, the 3D structure of brain organoids makes it difficult to combine with other approaches, such as MEA or electrophysiology in an undisturbed manner. The SNAR assay would be an excellent tool to monitor cell type-specific changes in brain organoids both for developmental profiling or in combination with pharmacology. Further, SNAR is compatible with human-iPSC derived neurons and thus should be easily applicable to studies in patient derived cell lines, which may harbor genetic defects otherwise difficult to recapitulate in animal models.

In addition, disease models often require intermingling of mutant and wild-type cells. This can now be achieved using CRISPR-Cas9 methods, but often leads to difficulties in the detection of a phenotype due to low signal to noise ratios (Aldinger et al., 2011; Sandoval et al., 2020). Given the sensitivity of the SNAR assay, these could now potentially be detected.

Overall, this simple live-cell assay will be useful to quantify and characterize neuronal responses upon genetic, pharmacological or other manipulations. The SNAR assay will be also applicable to large scale pharmacological screens and developmental profiling of patient iPSC-derived neurons.

